# Microbiota comparison of individual and pooled cow fecal samples from PEI dairy farms

**DOI:** 10.1101/2025.05.29.656927

**Authors:** Ana S. Jaramillo-Jaramillo, J T. McClure, Henrik Stryhn, Kapil Tahlan, Luke Heider, Javier Sanchez

**Affiliations:** Department of Health Management, Atlantic Veterinary College, University of Prince Edward Island, Charlottetown, Prince Edward Island, Canada; Department of Biology, Memorial University of Newfoundland, St. John’s, Newfoundland and Labrador, Canada

## Abstract

The aim of this study was to compare the microbiota diversity captured in individual versus pooled fecal samples from dairy cattle and evaluate the feasibility of using pooled sampling to assess the microbiota of dairy cattle at the herd level. A cross-sectional study used animals from 28 Prince Edward Island (PEI) dairy farms in Canada. The farms were visited between July and December 2020. Both free-stall and tie-stall housing systems were eligible. Manure samples from dry and lactating cows were obtained. Then, approximately 20g of fecal samples from each group were pooled. DNA extractions on all subsamples were performed using the Qiagen PowerMax Soil Kit and submitted for 16S rRNA gene amplification and sequencing. Operational taxonomic units were determined, and four alpha diversity indices were computed. A total of 128 and 132 manure samples from pos and prepartum cows, respectively, were analyzed. Mixed-effects random slope models were employed, incorporating herd-level random effects to estimate the correlation among individual samples and between individual and pooled samples. The estimated Shannon observed features, Pielou evenness and Faith PD were 5.9, 580, 0.93 and 44.1 from the individual and 5.93, 587, 0.93 and 45.5 from the pooled samples, respectively. All alpha diversity metrics were not significantly different between individual and pooled samples. Overall, pooled sampling does not significantly affect diversity and provides comparable results with individual samples, though it tends to show slightly higher diversity in some indices. This sampling strategy could be used in microbiota studies of dairy herds.

## Introduction

Metagenomics has emerged as a powerful tool for characterizing microbial communities, with fecal and intestinal microbiota sequencing providing key insights into the health, nutrition, physiology and environment of an animal (1). Microbiota study results are significantly influenced by sample collection, preservation, and processing methods, which can lead to considerable variation between individuals, even in similar environmental conditions. This high inter-individual variability in microbiota composition can introduce bias, increase costs, and reduce the number of sequence reads generated. These factors collectively have the potential to affect the biological interpretation of the data, posing challenges to consistent and reproducible findings (2,3).

In dairy farms, pooling fecal samples from multiple individuals within a herd has been suggested as a cost-effective and time-efficient strategy to assess overall or specific microbial abundance. This approach holds the potential for herd-level monitoring of health and antimicrobial resistance. However, the degree to which pooled samples accurately represent the microbial diversity found in individual samples remains uncertain, highlighting the need for further evaluation of its efficacy and limitations (4,5).

The aim of this study was to compare the microbiota diversity captured in individual versus pooled fecal samples from dairy cattle herds and evaluate whether pooled sampling can provide comparable insights at the herd level. To our knowledge, this is the first study to directly assess pooled versus individual fecal samples in dairy cattle herds, offering valuable insights into the feasibility and utility of pooled samples for herd-level microbiota analysis. We hypothesized that pooled sampling would accurately reflect herd-level microbial diversity, providing a practical and effective tool for health management, disease prevention, and antimicrobial resistance monitoring in dairy farming.

## Materials and methods

### Ethics statement

The owners of the selected farms provided permission to conduct this study. No additional permissions were required for the collection and analysis of samples.

### Study design and sample collection

For this observational cross-sectional study, 28 dairy farms from Prince Edward Island (PEI), Canada, participated in a research project focused on antimicrobial stewardship and resistance in dairy cattle. The farms were visited between July and December 2020. Inclusion criteria required a minimum herd size of 50 lactating cows, enrollment in the DHI (Herd Management Score) program, and the practice of raising their own heifers. Both free-stall and tie-stall housing systems were eligible.

### Sample selection and processing

A total of 137 individual fecal samples from the post and prepartum cows were collected from the 28 farms. Approximately 20 g of aliquots were taken from five cows. Each individual fecal sample was labelled with the cow’s ID and transported to the Atlantic Veterinary College (AVC) laboratory in coolers containing ice packs, ensuring the temperature remained below 8°C. To create the pooled sample, each individual sample was manually homogenized using a spatula to create one pooled sample, resulting in 28 pooled samples. Combined with the individual samples, this provided a total of 165 fecal samples from 28 farms. All samples were immediately stored in a –20°C freezer until DNA extraction.

### DNA extraction

DNA extractions on all subsamples were performed using the Qiagen PowerMax Soil Kit (Qiagen Laboratories) as per the manufacturer’s instructions. Frozen fecal samples were thawed at 4°C, and 0.25 grams of each subsample was used for the extraction. Estimated DNA yields and quality (A260/280 ratio) were verified using a NanoDrop 1000 Spectrophotometer (Thermo Fischer Scientific). A sample was re-extracted if it did not achieve a DNA yield of more than 30 ng/µL, a 260/280 ratio of over 1.80, and a 260/230 ratio of over 2.0. The extracted DNA was then stored at -80°C until it was shipped for sequencing.

### 16S rRNA gene sequencing and data processing

For microbiota sequencing, 100 µL of extracted DNA from each subsample was sent to the Integrated Microbiota Resource (IMR) laboratory (Dalhousie University, Halifax, NS, Canada) for 16S rRNA gene amplification and sequencing. The V6-V8 16S subunit bacteria-specific region was amplified with the primer set [5’-ACGCGHNRAACCTTACC-3’] / [5’- ACGGGCRGTGWGTRCAA-3’] to yield approximately 400-500 bp DNA fragments. Amplicon sequencing was performed on the Illumina MiSeq instrument (Illumina, San Diego, CA, USA) to produce 2 × 300 bp paired-end read lengths with up to 100,000 reads per sample. The sequenced 16S rRNA amplicons were analyzed using pipelines in Quantitative Insights Into Microbial Ecology version 2 (Qiime2-2020.11) tools (6). Briefly, all reads were processed for sequence quality and denoising using DADA2 (7); meanwhile, chloroplast and mitochondrial DNA sequences were removed from the dataset. All data and analysis files are available in our GitHub repository at the following link: XXX

### Taxonomy, diversity indices calculation and statistical analysis

Operational Taxonomic Units (OTUs) were assigned using a naive Bayes classifier trained on the Greengenes database (8) and were classified at phylum, class, order, family, and genus levels. Relative abundances of dominant bacterial populations were calculated for each sample type at the phylum and genus levels.

Alpha diversity metrics (Shannon, observed features, Pielou evenness, and Faith PD) of pooled and individual fecal samples were calculated at a rarefaction depth of 9400 seq/sample using the Phyloseq (9) and the DiversitySeq (10) packages in R version 4.0.4 (R Core Team, 2021). Mixed-effects random slopes models were employed, incorporating herd-level random effects and a structured correlation of sample type (pooled vs. individual) within the gestational status group (postpartum vs. prepartum). These models were used to estimate the correlations among the five individual fecal samples and between pooled and individual samples for each of the four alpha diversity indices. Standard errors for correlations were computed using the delta method. The correlations between sample type and standard errors were computed from the estimated variance components. The models included sample type, gestational status and their interaction (its cross-product term), and all terms were included regardless of their statistical significance. Before fitting the mixed models, linear models were fitted, and Box-Cox transformations were assessed to meet the assumptions of the regression models. All statistical analyses were performed with Stata 16.1 (StataCorp, College Station, Texas, USA) software. The figures were performed with the R package ggplot2 (version 4.0.4).

## Results

### Sample processing

Of the 165 DNA samples prepared in the current study, 160 were successfully sequenced, representing 28 dairy herds from 128 and 132 post and prepartum cows, respectively. Following sample trimming and alignment, an average of 19,855.7 reads per sample (range: 218–53,123) were obtained and analyzed. The individual sample group yielded an average of 19,488.3 reads per sample (range: 1,158–39,980), while the pooled sample group had an average of 21,588.3 reads per sample (range: 218–53,123).

### Taxonomy

The relative abundance of bacterial phyla in pooled and individual fecal samples is depicted in Figure 1. Firmicutes and Bacteroidetes dominate the microbial community in both sample types, collectively accounting for most of the abundance. Firmicutes appear to make up slightly more than 50% of the total abundance in both pooled and individual samples, while Bacteroidetes contribute around 30% to 35%. Other phyla, such as Actinobacteria and Verrucomicrobia, are present at much lower proportions. This distribution indicates a similar microbial composition at the phylum level across pooled and individual samples, supporting the potential of pooled samples to reflect the broader community structure observed in individual samples.

**Figure 1.**
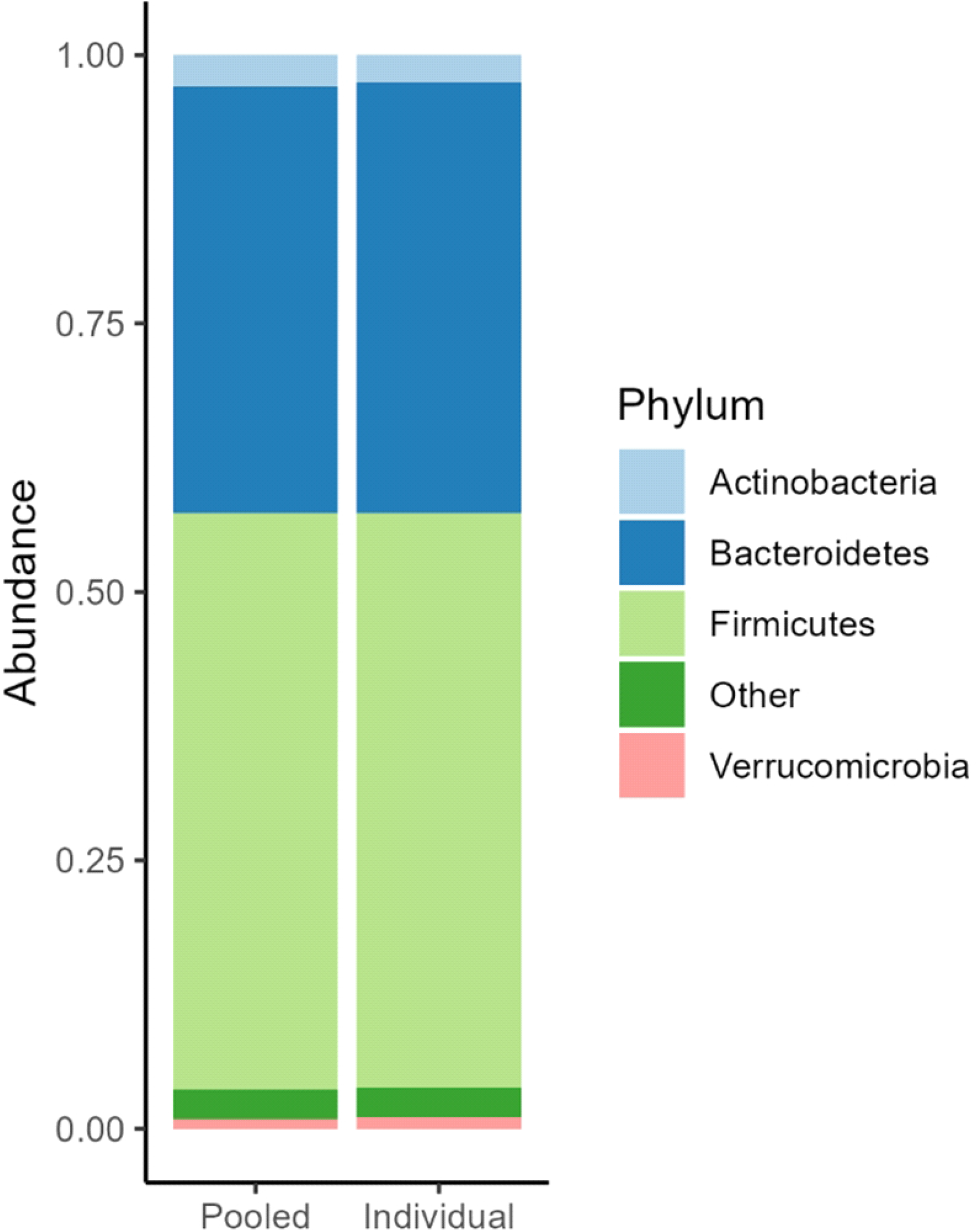
Bar plot of relative abundance at phylum level for two types of fecal samples from dairy cattle (Pooled and Individual)

Figure 2 illustrates the relative abundance of bacterial genera between pooled and individual fecal samples. In both sample types, “other genera” account for approximately 75% of the microbial community. Among the notable genera, Clostridium, 5-7N15, Akkermansia, and Bacteroides each contribute smaller proportions of the total abundance. Other genera, such as Ruminococcus, Turicibacter, and Prevotella, are present at lower levels. The overall distribution of genera is highly similar between pooled and individual samples, reinforcing the viability of pooled samples in representing the broader microbial community structure observed in individual samples.

**Figure 2.**
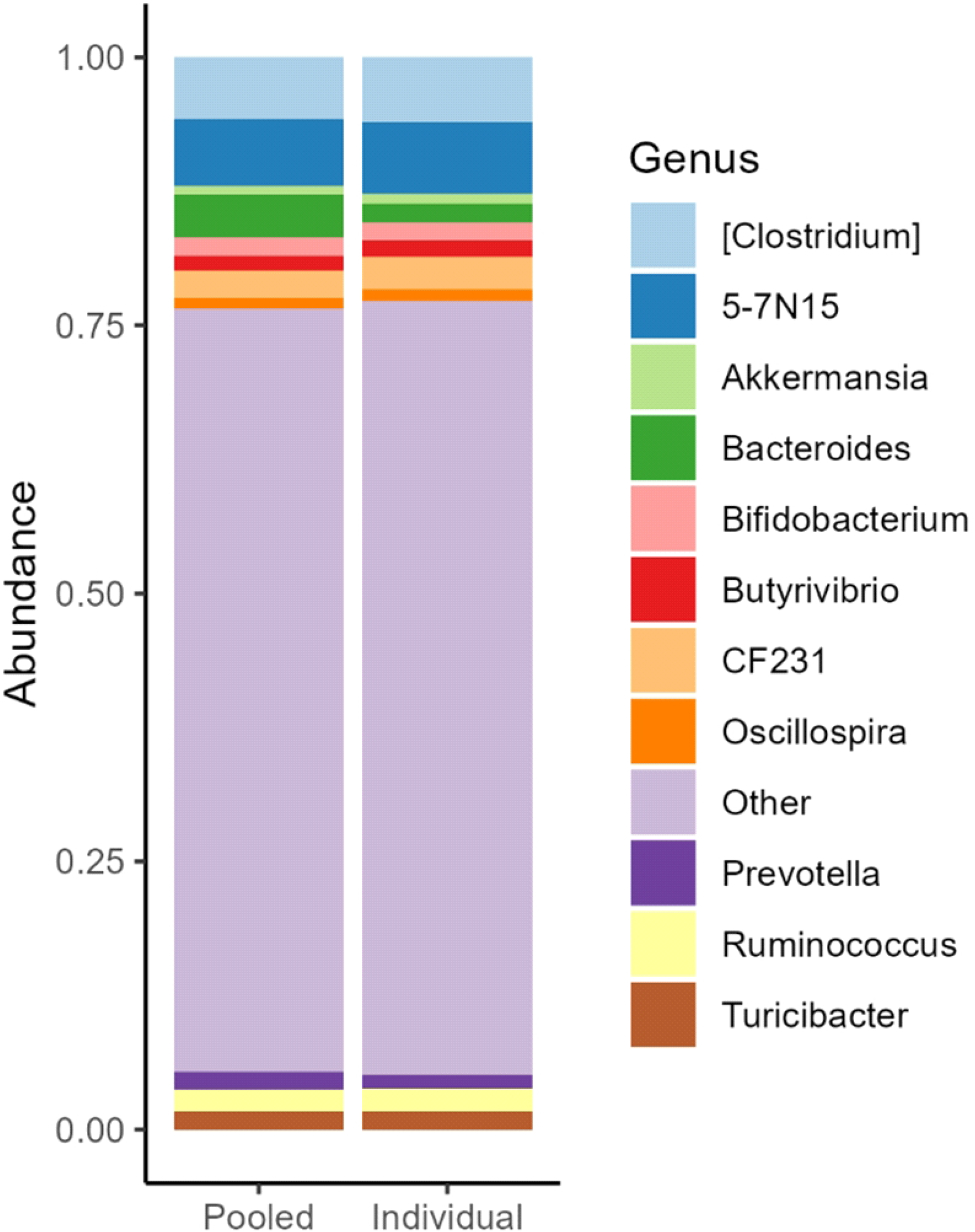
Bar plot of relative abundance at the genus level for two types of fecal samples from dairy cattle (Pooled and Individual).

### Alpha diversity

The means of all alpha diversity metrics were calculated for both individual and pooled groups. The estimated Shannon observed features, Pielou evenness and Faith PD were 5.9, 580, 0.93 and 44.1 from the individual and 5.93, 587, 0.93 and 45.5 from the pooled samples, respectively. The distribution of the values for each of the four indices evaluated is depicted in Figure 3.

**Figure 3.**
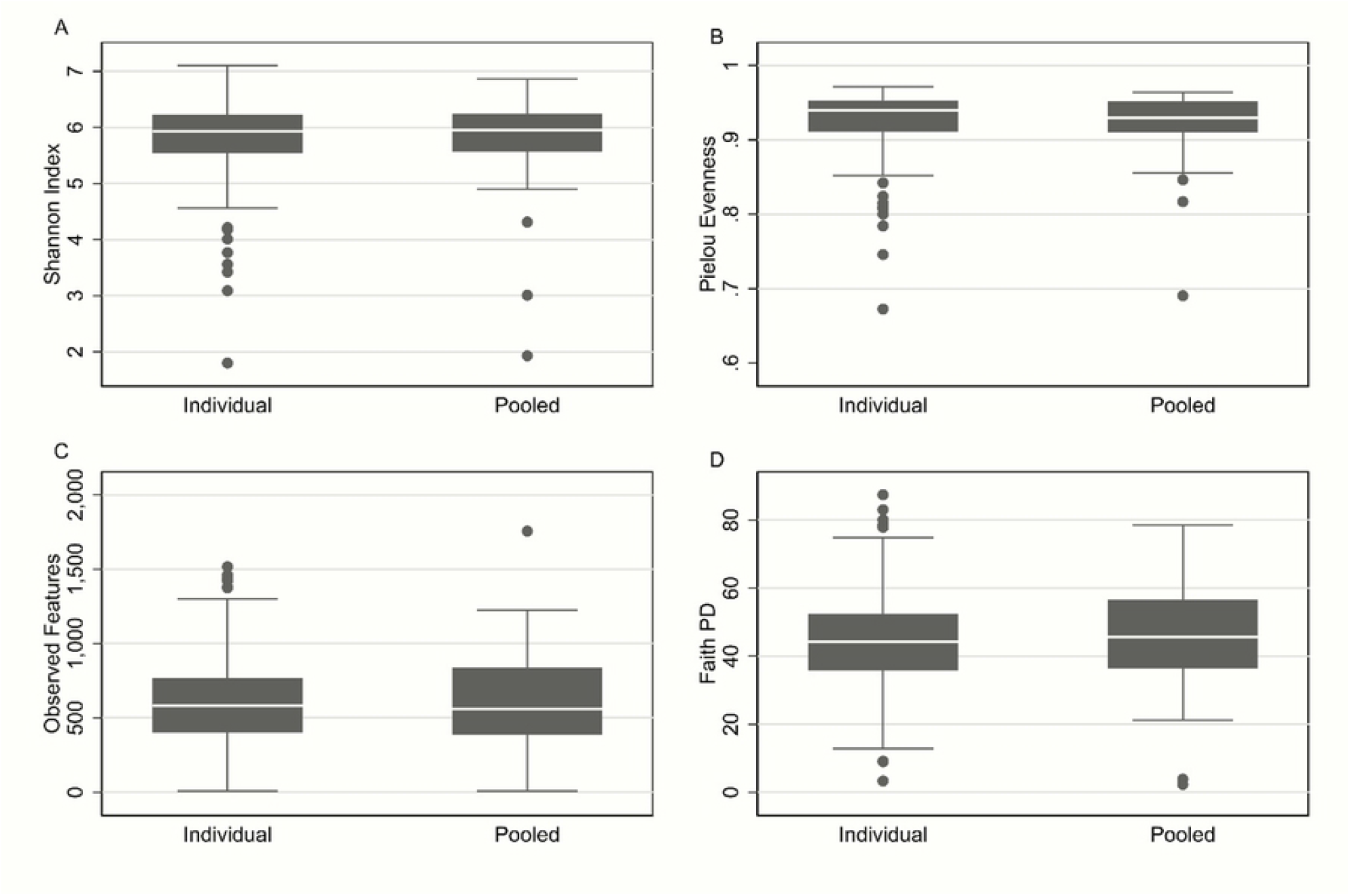
Box plots of alpha diversity indices (A) Shannon, (B) Pielou Evenness, (C) Observed features, and (D) Faith PD, measured for individual and pooled fecal sample types. For better visualization, the y-axis scale in plot B was adjusted.

The mixed-effects random slopes models evaluating the impact of sample type and gestational status on the four alpha diversity indices indicated a non-significant interaction between sample type and gestational status and all the indices indicated a significant decrease in diversity in postpartum samples as compared to samples taken in the dry period (e.g. prepartum) (Table 1). Additionally, all the models indicated an important variability in microbiota diversity across herds.

**Table 1:**
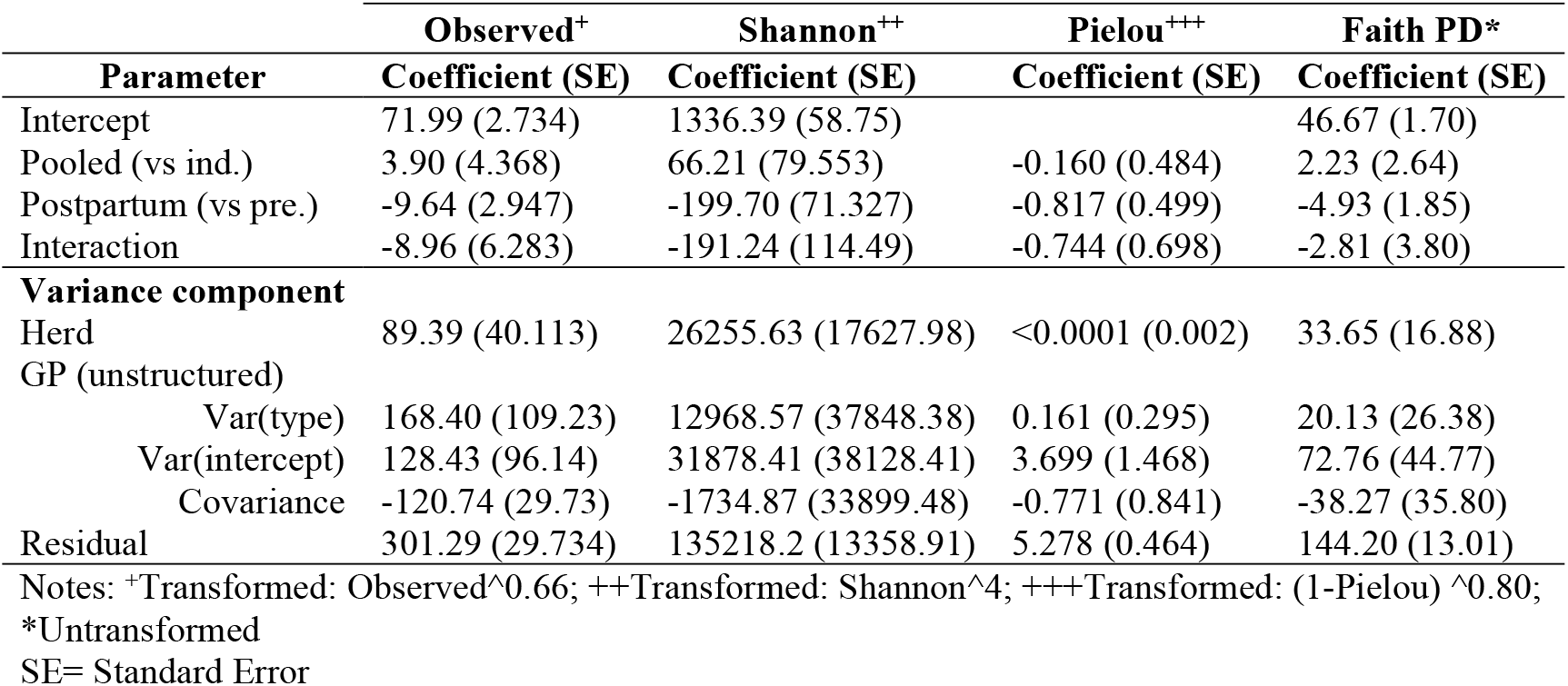
Mixed effects random slope model coefficients and variance components for each of the four alpha diversity indices analyzed from 154 and 160 post and prepartum dairy cows, respectively, from 28 herds.

The estimated correlations [and standard errors] of alpha diversity indices between individual and pooled samples are summarized in Table 2. Overall, the correlations were small but non-significantly different between sample types (e.g. individual vs pooled).

**Table 2.** Estimated correlations and standard errors between individual and pooled samples for each of the alpha diversity indices.

|  | Individual - Individual | Individual - Pooled |
| --- | --- | --- |
| Index | Rho (SE) | Rho (SE) |
| Observed | 0.324 (0.071) | 0.202 (0.093) |
| Shannon | 0.333 (0.069) | 0.285 (0.056) |
| Pielou | 0.305 (0.064) | 0.355 (0.064) |
| Faith PD | 0.257 (0.069) | 0.309 (0.068) |

Notes: SE= Standard Error; Rho: Correlation

## Discussion

Overall, pooled sampling does not significantly affect diversity, though it tends to show slightly higher diversity in some indices. Pooled fecal samples were shown to effectively represent the microbial diversity information reflected in alpha diversity indices, offering a cost-effective option for herd-level microbiota studies in dairy cattle. However, if the objective is to quantify the absolute abundance or richness of a specific OTU, phyla or genera or to assess their proportions at the herd level, pooled samples may not be ideal, as several individual samples were shown to capture more detailed information. In contrast, other alpha diversity metrics like Shannon and Pielou’s evenness are similarly represented by both pooled and individual samples, supporting the use of pooled samples for these indices.

For observed diversity, the lower correlation between individual-pooled samples suggests that pooling may reduce the precision of diversity estimates. This is likely due to the combined effect of multiple individuals contributing to a single pooled measurement, which may dilute the distinct microbial signatures of individual samples. On the other hand, Shannon diversity shows only a modest correlation reduction, implying that pooled samples can still provide reasonable estimates for this index. Pielou’s evenness demonstrates a slightly higher correlation between individual-pooled samples, indicating that pooling does not significantly distort evenness estimates. For Faith PD, the higher correlation between individual and pooled samples suggests that pooling may be advantageous in capturing phylogenetic diversity, however the differences with the correlations between individual samples were minimal and non-statistically significant. This is likely because pooling aggregates broader evolutionary lineages, offering a more comprehensive representation of the microbial phylogeny at the herd level.

In the case of phenotypic diversity, indices like Faith PD measure the diversity of a subset of taxa by summing the lengths of the branches on the phylogenetic tree connecting all taxa within the subset (11). The correlation between individual samples was the weakest; however, not significantly different from the other indices. However, for this index, this could be due to the influence of unique individual microbial communities and the rapid changes these communities can undergo over a short period, even under similar environmental conditions. However, the correlation between pooled and individual samples was similar, suggesting that pooled samples can adequately represent the overall fecal microbial structure of the herd. This highlights the potential utility of pooled sampling as a practical and cost-effective approach for herd-level microbiota analysis.

The significant herd-level variation observed in all the models suggests that diversity patterns are influenced by herd-specific factors, such as management practices, environmental conditions, or host genetics, which are not accounted for by the fixed effects in the models. The analysis reported in Table 2 shows that correlations between individual and pooled samples vary by diversity index, with Pielou’s evenness and Faith PD showing greater consistency.. Correlations between individual samples are generally moderate across all indices, indicating consistency in diversity estimates in such cases. However, the correlations between individual and pooled samples display more variability but are still very similar to those between individual samples. This variability underscores the nuanced trade-offs of using pooled samples for herd- level diversity assessments.

Our findings suggest that deciding to pool samples versus analyzing them individually may have different implications depending on the studied diversity index. While pooling might introduce some variation, particularly for observed diversity, it provides more consistent estimates for indices like Pielou’s evenness and Faith PD. Therefore, the choice of sampling method should consider the specific diversity indices of interest and the study’s goals. For taxonomy, the relative abundance profiles at the phylum and genus level in pooled and individual samples are similar, suggesting that pooling of samples does not drastically alter the overall microbial composition. This indicates that pooled samples may be a viable approach for capturing the broader community structure observed in individual samples, making them an option for studies focused on general microbial community dynamics. In addition, our results underscore the influence of gestational status and herd-specific factors on diversity metrics, highlighting the complexity of microbiota studies.

Given the distinct characteristics exhibited by the microbiota of pooled samples, they preserved key microbial signatures observed in individual samples.. However, the choice of sampling technique for assessing the fecal microbiota within a dairy farm should align with the specific objectives of the study being conducted. The microbiota profile of a single individual may fluctuate due to various factors. Thus, multiple individual samples offer a more comprehensive herd perspective than a pooled sample from the same population. However, sequencing the microbiota of pooled samples could prove valuable in characterizing the overall microbial community within the herd. This approach may also provide crucial insights into the roles of specific taxonomic features under varying conditions or treatments, making it a useful strategy for certain herd-level investigations.

